# The Effects of Pitch Shifts on Delay-induced Changes in Vocal Sequencing in a Songbird

**DOI:** 10.1101/072009

**Authors:** MacKenzie Wyatt, Emily A. Berthiaume, Conor W. Kelly, Samuel J. Sober

## Abstract

Like human speech, vocal behavior in songbirds depends critically on auditory feedback. In both humans and songbirds, vocal skills are acquired by a process of imitation whereby current vocal production is compared to an acoustic target. Similarly, performance in adulthood relies strongly on auditory feedback, and online manipulations of auditory signals can dramatically alter acoustic production even after vocalizations have been well learned. Artificially delaying auditory feedback can disrupt both speech and birdsong, and internal delays in auditory feedback have been hypothesized as a cause of vocal dysfluency in persons who stutter. Furthermore, in both song and speech online shifts of the pitch (fundamental frequency) of auditory feedback lead to compensatory changes in vocal pitch for small perturbations, but larger pitch shifts produce smaller changes in vocal output. Intriguingly, large pitch shifts can partially restore normal speech in some dysfluent speakers, suggesting that the effects of auditory feedback delays might be ameliorated by online pitch manipulations. While birdsong provides a promising model system for understanding speech production, the interaction between sensory feedback delays and pitch shifts have not yet been assessed in songbirds. To investigate this, we asked whether the addition of a pitch shift modulates delay-induced changes in Bengalese finch song, hypothesizing that pitch shifts would reduce the effects of feedback delays. Compared the effects of delays alone, combined delays and pitch shifts resulted in a significant reduction in behavioral changes in one type of sequencing (branch points) but not another (distribution of repeated syllables).

**Significance Statement:** Vocal behavior depends critically on an organism’s ability to monitor the sound of its own voice (“auditory feedback”). Studies of both humans and songbirds have demonstrated that successful vocal performance depends critically on the quality and timing of such feedback, however the interaction between vocal acoustics and the timing of auditory feedback is unclear. Here we used songbirds to examine this interaction by measuring vocal performance during delays and distortions (pitch shifts) of auditory feedback.

## Introduction

Learned vocal behaviors depend strongly on auditory feedback. In both birdsong and human speech, adults rely on auditory feedback to detect and correct errors in vocal production. This reliance on auditory information can be demonstrated by manipulating auditory feedback and measuring the effects on vocal output.

Complete elimination of auditory feedback by and deafening in adulthood leads to dramatic vocal performance deficits (McGarr, 1983; Okanoya and Yamaguchi, 1997; Woolley and Rubel, 1997; Lombardino and Nottebohm, 2000). More subtle manipulations of auditory signals reveal the complex influence of sensory feedback on motor programming. Artificially delaying auditory feedback in human speakers can cause vocal sequencing errors, including unwanted repetitions of consonants and words, in normally fluent speakers (Fairbanks, 1955; Chase, 1958; Yates, 1963). Such results suggest that the sequencing errors observed in persons who stutter might result from disorders of auditory feedback processing (Buchel and Sommer, 2004; Hampton and Weber-Fox, 2008). Intriguingly, artificially delaying auditory feedback is sometimes effective as a treatment for stuttering (Ryan and Van Kirk, 1974; Kalinowski and Stuart, 1996), further linking the dependence of vocal sequencing on the timing of auditory feedback and emphasizing the complex relationship between sensory feedback and speech production. Analogously, studies of birdsong have shown that perturbations of auditory feedback timing can degrade vocal production. Delayed playbacks of a bird’s own syllable during singing leads to song degradation after chronic exposure in zebra finches (Leonardo and Konishi, 1999; Cynx and von Rad, 2001). In Bengalese finches, a species whose song contains “branch points” where vocal sequencing is probabilistic rather than fixed (Fig. 1), acute changes in vocal sequencing can result from delayed playbacks of a bird’s own song syllable while singing (Sakata and Brainard, 2006). The similarities of these results across species suggest songbirds as a promising animal model for disorders of human speech production.

**Figure 1:**
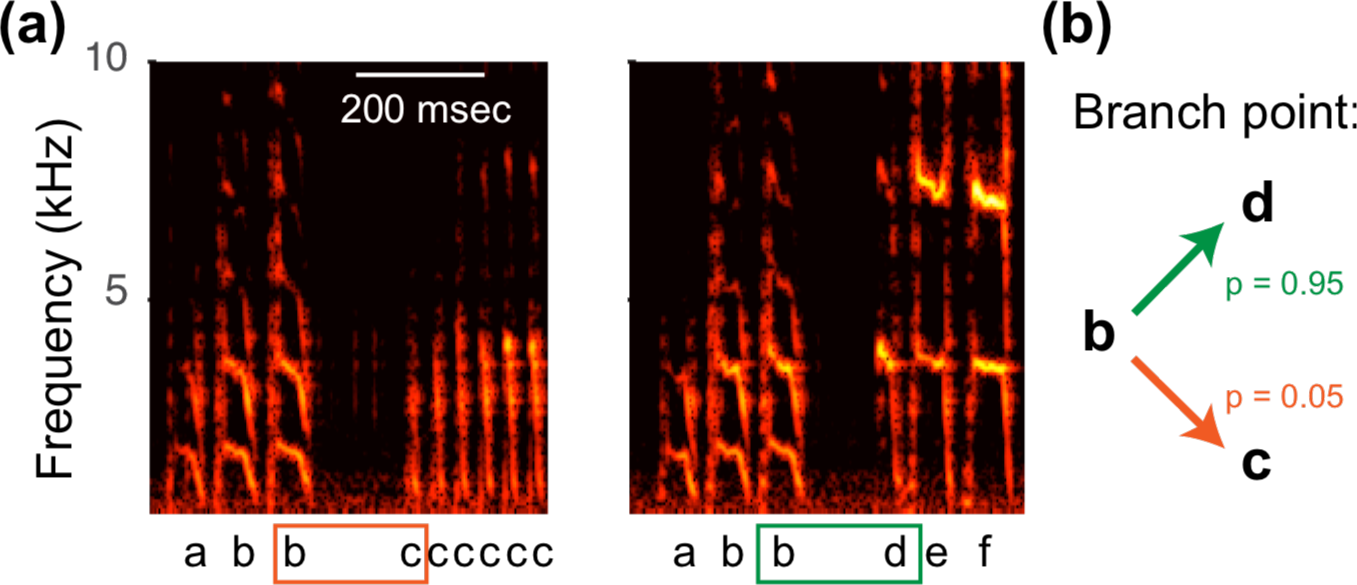
Sequence variation in birdsong. **(a)** Spectrographic representations show the power (heat map) at each acoustic frequency (vertical axis) as a function of time (horizontal axis). The two spectrograms show excerpts from different times during the same bout of song from a single bird. Labels below the spectrogram indicate different syllables. Orange and green boxes highlight a “branch point” in which syllable “b” can be followed by either syllable “d” or syllable “c”. **(b)** Schematic quantifies transition probabilities for this branch point.

Other studies in songbirds and humans have explored how the brain uses the acoustic structure of auditory feedback (as distinct from the timing of feedback) to calibrate vocal performance. In both songbirds and humans, manipulations of the fundamental frequency (which we will refer to here as “pitch”) of auditory feedback evoke compensatory responses, for example by increasing the pitch of vocal output in response to a decrease in the pitch of online auditory feedback (Jones and Munhall, 2000; Sober and Brainard, 2012; Hoffmann and Sober, 2014). Notably, vocal pitch changes in birdsong and formant changes in human speech are most robust for smaller shifts in auditory feedback, with larger shifts evoking little or no change in vocal output (Burnett et al., 1998; Liu and Larson, 2007; MacDonald et al., 2010; Katseff et al., 2012; Sober and Brainard, 2012), suggesting that during large pitch shifts, the brain relies less on auditory feedback to influence ongoing vocal behavior. Intriguingly, large (half an octave) pitch shifts cause an increase in speech fluency in some persons who stutter (Kalinowski et al. 1993; Natke et al. 2000), suggesting an interplay between the acoustic structure of auditory feedback and the sequencing of vocal motor commands. However, the relationship between the acoustics and timing of vocal feedback, and their influence on vocal output, remain poorly understood.

We used Bengalese finches to investigate whether delay-induced changes in vocal production were influenced by alterations in the pitch of auditory feedback. Although the effects of auditory feedback delays on song have been tested previously using song-triggered playbacks of previously-recorded songs, technical challenges have prevented the use of continuous delayed feedback. We overcame this obstacle using miniaturized headphones (see Methods), which provided continuous, delayed feedback in real time (Hoffmann et al., 2012). Experimental conditions included a null condition (no delay or pitch manipulation), delayed auditory feedback (DAF) without any pitch shift, and a condition in which auditory feedback was both delayed and pitch shifted (DAF+PS). We predicted that, in agreement with prior findings using playbacks of short segments of song (Sakata and Brainard, 2006), delayed feedback would induce changes in the syllable transition probabilities. In particular, we hypothesized that, as reported by Sakata and Brainard (2006), delayed feedback would result in the most common transition (the “primary” transition) becoming less prevalent, and the non-primary transition becoming more common. We further hypothesized that the large pitch shift in the DAF+PS condition would reduce the magnitude of the changes observed in the DAF condition.

## Materials and Methods

Four adult (>100 days old) male Bengalese finches (*Lonchura striata* var. *domestica*) were used as the experimental subjects. During experiments, birds were housed individually in an isolated sound-attenuating chamber, and all song was undirected (i.e. produced in the absence of female birds). The light/dark cycle was maintained for 14 h:10 h, with lights on beginning at 7 AM and ending at 9 PM. All procedures were approved by [Author University] Institutional Animal Care and Use Committee. *Experimental procedure.*

Miniature, lightweight headphones were custom-built out of lightweight carbon fiber and custom-fit to each bird’s head (Hoffmann et al., 2012). A condenser microphone in the bird’s cage (Fig. 2a) captured the birds acoustic output, which was routed to online sound-processing hardware (Eventide H7600), which provided perturbations of the pitch and/or timing of auditory feedback in real time. This manipulated feedback was then relayed to miniaturized speakers (EH-7157-000, Knowles) inside the headphones. In addition to the speakers, the headphones apparatus included a miniaturized microphone (EM-3046, Knowles) placed between the speaker and the opening of one of the ear canals. This microphone allowed us to monitor the performance of the headphones apparatus. The miniaturized microphone was used to calibrate the system such that the acoustic signal played through the headphones speakers was ~2 log units greater than auditory feedback leaking through the carbon fiber frame. The headphones therefore shielded the birds’ airborne vocalizations, allowing the altered feedback to replace the natural version. As described in detail below, auditory feedback conditions included a null condition in which no manipulation was introduced, a “delayed auditory feedback” (DAF) condition in which auditory feedback was delayed by 175 msec, or a “delay + pitch shift” (DAF+PS) condition in which both a 175 msec delay and an upward or downward pitch shift were applied simultaneously. As detailed previously, the sound processing hardware relayed the online acoustic signal to the headphones with a minimal delay (i.e. when auditory feedback was not being intentionally delayed) of ~10 msec, a delay which does not in itself evoke any measurable changes in vocal behavior (Sober and Brainard, 2009; Hoffmann et al., 2012; Kelly and Sober, 2014).

**Figure 2:**
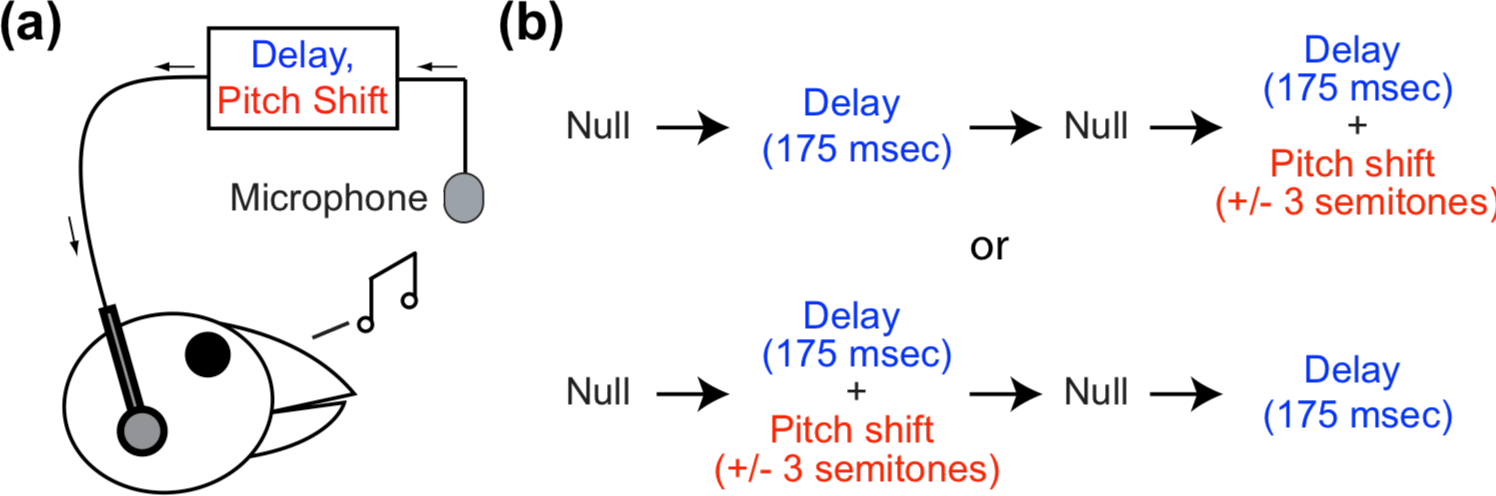
Experimental design. **a.** Auditory feedback manipulated using miniaturized headphones. A microphone transmits a bird’s vocalizations to online sound processing hardware, which are used to introduce a 175-msec delay (with or without a ±3 semitone pitch shift). **b.** Schedule of experimental conditions. Each bird was exposed to the delayed feedback alone (DAF) and delayed feedback plus a pitch shift (DAF+PS). The order of these conditions, as well as the direction of the pitch shift, was randomized across subjects. Prior to each DAF or DAF+PS epoch, birds sang in a “null” epoch free of delays or pitch shifts. The four experimental epochs lasted five days each.

The experimental design is outlined in Figure 2b. First, once birds habituated to the headphones, all birds sang during a period with zero pitch shift or delay for five days. After this null period, in the example shown at top in Figure 2b, the birds’ auditory output was altered with delayed auditory feedback (“DAF block”) for five days. The birds heard their natural vocalizations at 175 msec delay relative to output. We selected this delay magnitude after preliminary studies in two birds (not used in the present study) suggested that this delay consistently evokes changes in vocal sequence; however different delay values were not tested systematically. After the altered auditory feedback block, birds were subjected to second five-day null period of singing with zero pitch shift and no introduced delay. The birds were then subjected to a delayed auditory feedback and pitch shift block (“DAF+PS”) lasting five days. During the DAF+PS block, delayed feedback at 175 msec was concurrently pitch shifted up or down three semitones. Both the sign of the pitch shift (i.e. upward or downward by 3 semitones) in the DAF+PS condition and the order of the DAF/DAF+PS blocks were varied across birds to counterbalance for any learning order effects.

### Measuring song syntax features

As in previous uses of the headphones paradigm (Sober and Brainard, 2009), we analyzed songs produced during a fixed time window (here, 8 am - 12 noon). In cases where birds produced more than sixty bouts of song during this interval, we used only sixty bouts (spaced evenly across the interval) in the analysis. Syllable onsets and offsets were determined using an amplitude threshold, and song syllables were assigned arbitrary labels (e.g. a-f in Fig. 1) by visual inspection. Note that the use of the same letters for labeling syllables across different birds does not indicate acoustic similarities between the birds’ syllables.

We examined syllable sequencing in two contexts: branch points and repeated syllables. At a branch point, a single syllable can be followed by multiple different syllables. Such sequence variability is a hallmark of Bengalese finch song (Okanoya, 2004; Wohlgemuth et al., 2010; Matheson and Sakata, 2015), and branch point probabilities are actively maintained during vocal learning (Warren et al., 2012). At each branch point we quantified the probability of each transition (e.g. Fig. 1b). We used a z-test for proportions to compare probabilities from altered auditory feedback conditions to the null condition immediately preceding it. For a group analysis of the effects of DAF and DAF+PS on branch point probabilities across birds, we used a one-sided Wilcoxon signed rank test to evaluate our hypothesis that the effects of delayed auditory feedback would be reduced if the delay were performed in the presence of a pitch shift. We performed this statistical test only on changes in the probability of the primary (most common) transition, for two reasons. First and most importantly, the probabilities of primary and non-primary transitions at a single branch point are not independent. For example, if there are only two transitions and one increases by 10%, then the other must decrease by the same amount, so it would be incorrect to consider changes in two transitions at a single branch points as separate measurements. Second, we focused on the primary transition to evaluate our hypothesis that delayed auditory feedback (in both the DAF and DAF+PS condition) would lead to a reduction in the probability of the primary transition, as observed previously in a similar experiment (Sakata and Brainard, 2006). In total, the four birds used in our studies yielded a total of 9 branch points (1-4 per bird), consisting of four cases in which one syllable could be followed by one of two different syllables, and five cases in which one syllable could be followed by one of three different syllables.

Bengalese finches also commonly produce “repeated” syllables (e.g. syllable “c” in Fig. 3a), which are produced multiple times in succession. We quantified the distribution of repeat numbers for each repeated syllable in each tested auditory feedback condition (for example, the excerpt of song shown in Fig. 3a contains a case in which syllable “c” is repeated four times). We used a Kolmogorov-Smirnov test to determine whether the repeat distributions of individual syllables differed significantly across feedback condition, comparing the repeat distribution in the DAF or DAF+PS condition with that of the null period immediately preceding it. As in the analysis of branch point probabilities, we used a Wilcoxon signed-rank test to determine whether the change from null to DAF was significantly larger than the changes induced by the DAF+PS. In all statistical tests of both branch points and repeated syllables, we used data from only the last three days of each auditory feedback conditions. The four birds examined yielded a total of 20 repeated syllables (3-9 per bird). In all analyses described above, for each branch point or repeated syllable we combined data across the last three days of the five-day feedback epoch (null, DAF, or DAF+PS) when computing the effects of each feedback condition.

**Figure 3:**
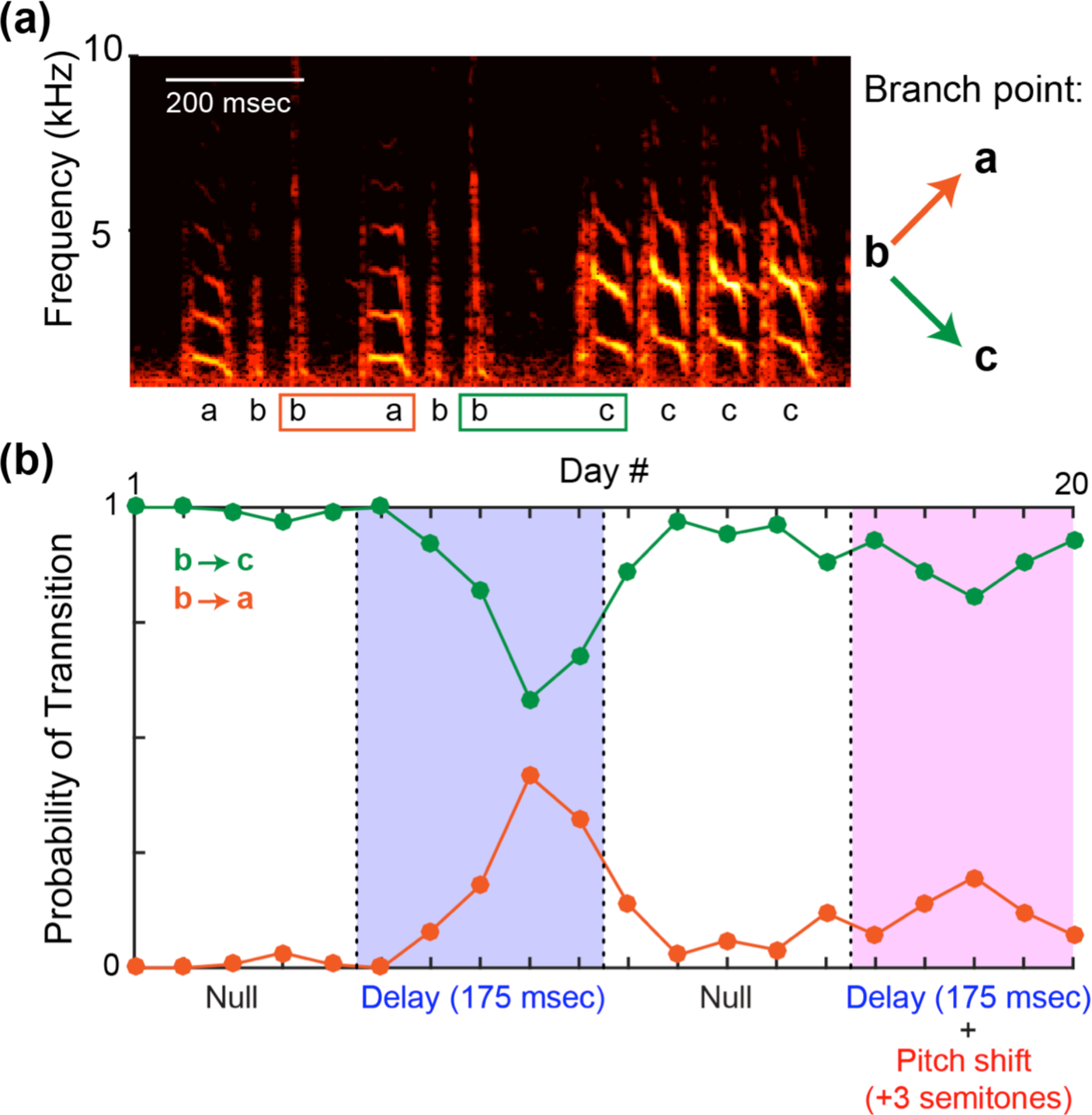
Effects of auditory feedback manipulations on branch point probabilities. **(a)** One branch point (in which syllable “b” can be followed by either syllable “a” or “c”) in our dataset. Spectrogram plotting conventions as in Figure 1a. **(b)** In this experiment, after a null period of unmanipulated auditory feedback the bird experienced delayed auditory feedback (DAF), followed by another period of unmanipulated feedback, followed by a combined delay and pitch shift (DAF+PS). Green and orange traces show the probability of the b➔c and b➔a transitions, respectively.

## Results

As hypothesized, delaying auditory feedback often induced changes in syllable sequencing. Figure 3 shows data from one branch point. In the first null period, syllable “b” was followed by syllable “c” more than 95% of the time (green trace, Fig. 3b), and was therefore the primary transition at that branch point (see Methods). Syllable “b” was followed by syllable “a” less than 5% of the time (orange trace, Fig. 3b). During the DAF condition (blue shaded region, Fig. 3b) transition probabilities gradually shifted, with the b➔c transition becoming less common and the b➔a transition becoming more common. Figure 4a summarizes the effects of DAF on transition probabilities across all branch points examined. In 7 of 9 cases, delayed feedback led to a significant reduction in the probability of the primary transition (Fig. 4a, filled symbols, p<0.05, z-test for proportions). When considered as a group, transition probabilities decreased significantly as a result of DAF being applied (p<0.01, one-sided Wilcoxon signed rank test).

**Figure 4:**
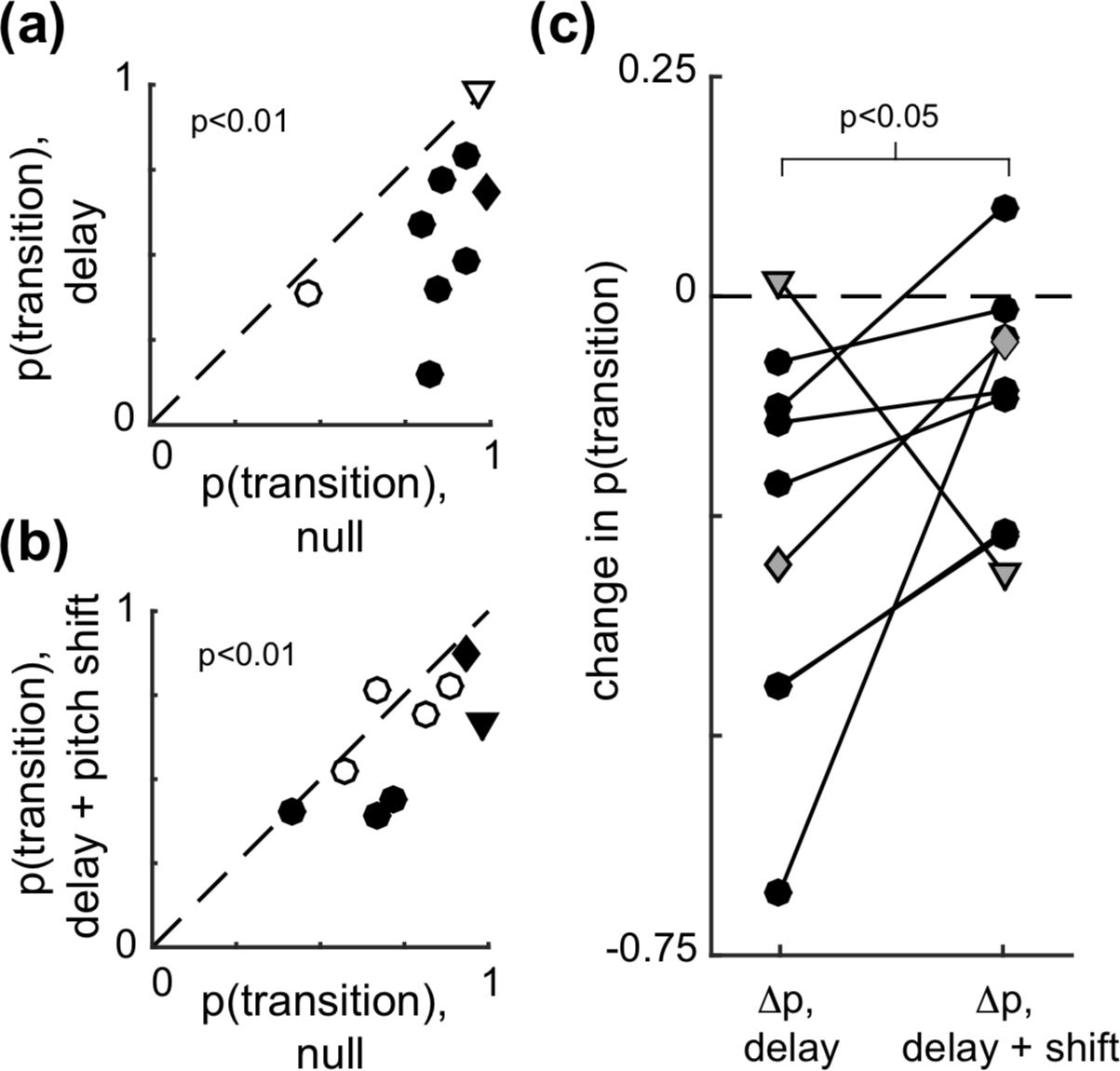
Effects of auditory feedback manipulations on branch points (group data). **(a)** Probability of the primary (i.e. most common, see Methods) transition in the null versus DAF conditions. Filled circles indicate that the difference between each probability in the null and DAF condition was statistically significant (p<0.05, z-test for proportions). Across all cases, transition probabilities significantly decreased as a result of DAF (p<0.01, one-sided Wilcoxon signed rank test) **(b)** Transition probabilities in the null versus DAF+PS condition. Other plotting conventions as in (a). **(c)** Comparison of the change in transition probability induced by the DAF (“delay”) and DAF+PS (“delay + shift”) conditions. As hypothesized, the change in probability was significantly smaller in the DAF+PS condition (p<0.05, one-sided Wilcoxon signed rank test). In all plots, diamond symbols indicate data from the example shown in Figure 3, triangle symbols indicate data from the example shown in Figure 5.

In the example shown in Figure 3, the DAF+PS condition (pink shaded region) had a similar, but smaller, effect on syllable sequencing than DAF, with the b➔c transition becoming slightly less prevalent in the DAF+PS epoch compared to the preceding null period. Figure 4b shows the effects of DAF+PS across transition points. Similar to the DAF condition (Fig. 4a), DAF+PS induced significant changes in most cases (Fig 4b, filled symbols, p<0.05, z-test for proportions) and as a group exhibited a significant reduction in the probability of the primary transition (p<0.01, one-sided Wilcoxon signed rank test).

We then asked whether, consistent with our hypothesis, the changes induced by DAF+PS were smaller than those induced by DAF alone. Figure 4c compares the change in transition probability induced by DAF (“Δp, delay”) with that induced by DAF+PS (“Δp, delay + shift”). As hypothesized, the effects of DAF+PS were significantly smaller (p<0.05, one-sided Wilcoxon signed rank test).

Notably, although overall DAF+PS produced significantly smaller changes in transition probability than DAF, in one case (triangle symbols in Fig. 4), much larger changes were observed in the DAF+PS condition. Data from this branch point is shown in Figure 5. Therefore, it is important to emphasize that although the group analysis demonstrated smaller changes once pitch shifts were added to delays, the opposite was seen in one individual case.

**Figure 5:**
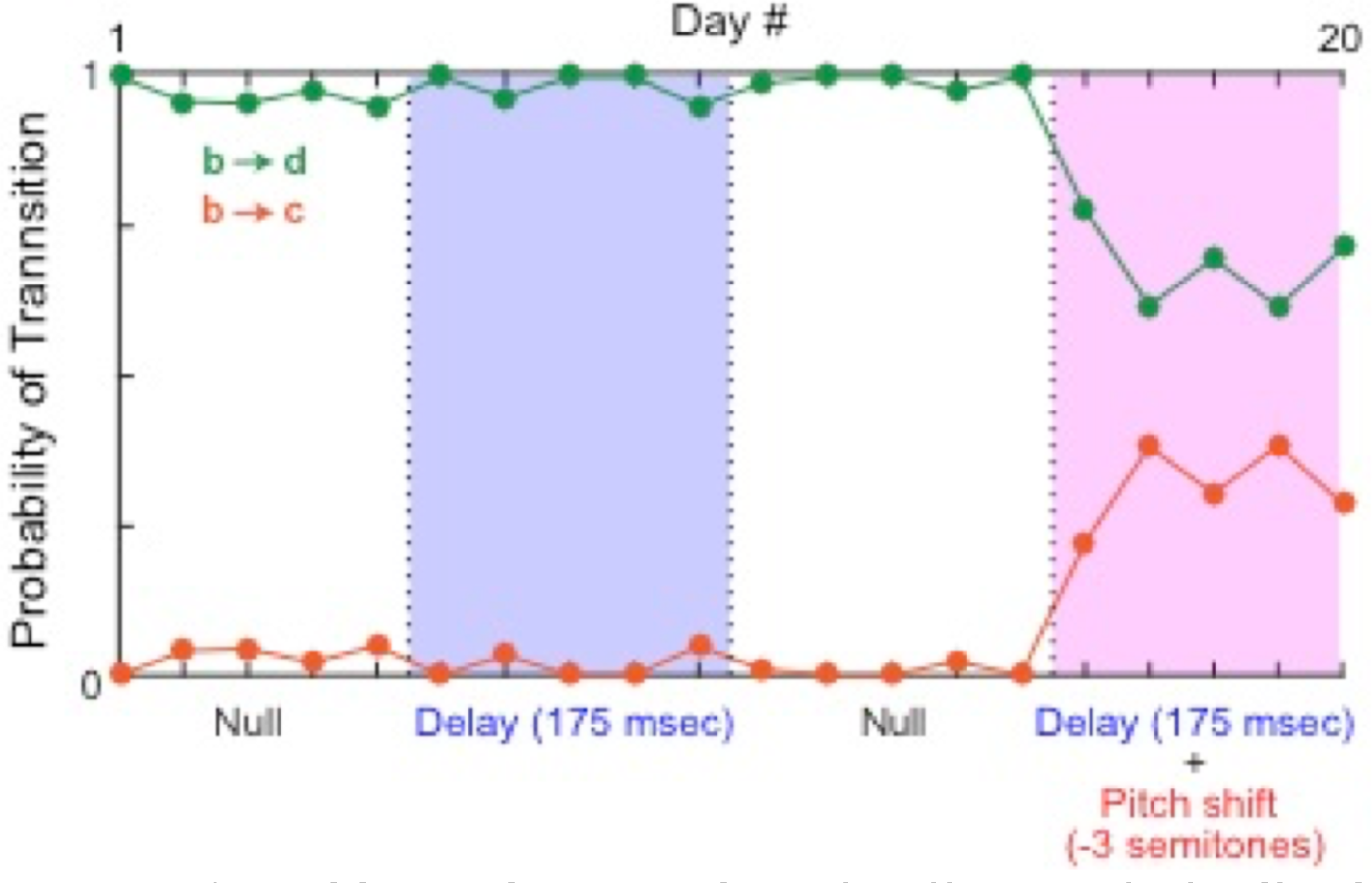
Additional example of effects of feedback manipulations on branch point probabilities. Plotting conventions as in Figure 3. Data are from the branch point also shown in Figure 1.

We also examined the effect of auditory feedback manipulations on the distribution of repeated syllables (Fig. 6a shows an example containing four different repeated syllables). Figure 6b shows an example from our dataset in which DAF induces a significant change in the distribution of repeats of syllable “g” (p<0.05, Kolmogorov-Smirnov test). As shown in Figure 7a, DAF frequently led to significant changes in repeat distribution (filled symbols), although there was no significant bias towards increases or decreases in mean repeat number (p=0.13, two-sided Wilcoxon signed rank test). The DAF+PS condition (Fig. 7b) also induced significant changes in the repeat distribution in many cases. Interestingly, in the majority (16/20) of these cases, DAF+PS reduced the mean number of repeats (p<0.01, two-sided Wilcoxon signed rank test). Figure 7c and d show the same data as Figure 7a and b, respectively, but represented as a change in repeat number between the null and DAF/DAF+PS conditions.

**Figure 6:**
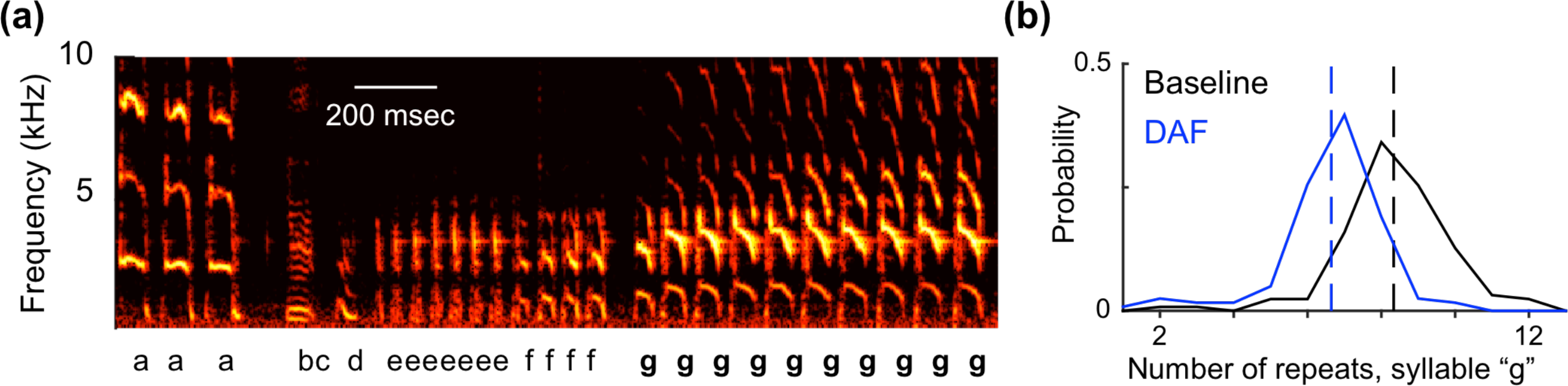
Effects of auditory feedback manipulations on repeat lengths. **(a)** Spectrogram shows excerpt of a song that contains three repeated syllables (“e”, “f”, and “g”). In this excerpt, syllable “g” is repeated 10 times. Spectrogram plotting conventions as in Figure 1a. **(b)** Distribution of repeat lengths of syllable “g” in the null condition (black solid line) and DAF condition (blue solid line). Dashed lines show the mean of each distribution.

We next evaluated our hypothesis that the DAF+PS condition would evoke smaller changes in repeat number than the DAF condition. Comparing these changes (Fig. 7e) did not reveal any significant difference between the two conditions (p=0.65, two-sided Wilcoxon signed rank test). We futher asked whether any differences existed between the data from the two conditions shown in Fig. 7e by performing a two-sample Kolmogorov-Smirnov test, which similarly failed to detect any significant difference (p=0.77). Therefore, although in individual cases both branch point probabilities (Fig. 4a, b) and repeat number (Fig. 7a, b) were often significantly modulated by DAF and/or DAF+PS, the effects of these two alterations of auditory feedback differed significantly only for branch point probabilities (Fig. 4c) but not for the distribution of repeated syllables (Fig. 7e).

**Figure 7:**
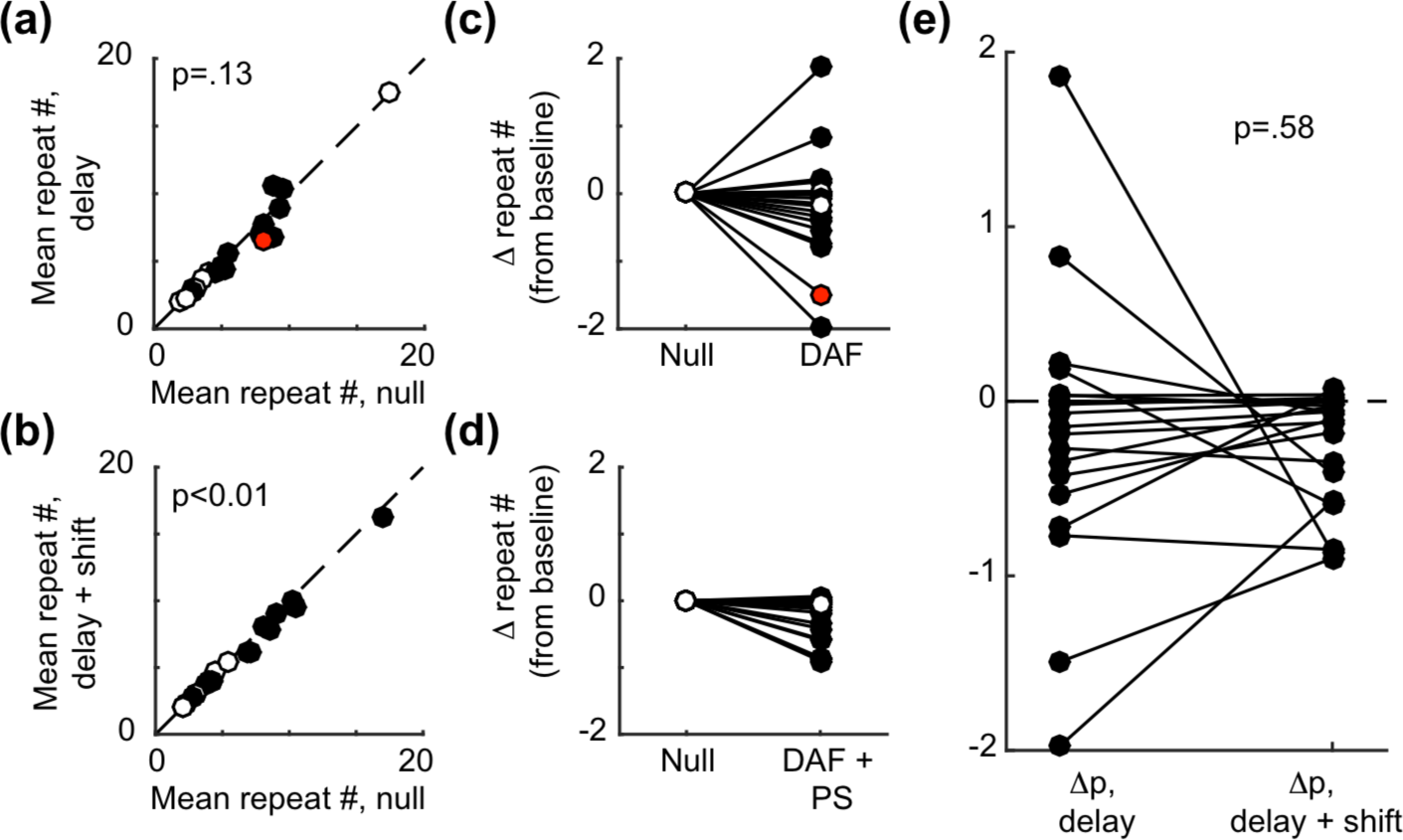
Effects of auditory feedback manipulations on repeat lengths (group data). **(a)** Mean repeat length in the null versus DAF conditions. Filled circles indicate that the difference between the repeat distribution in the null and DAF condition was statistically significant (p<0.05, two-sample Kolmogorov-Smirnov test). Across all cases, mean repeat numbers did not differ significantly as a result of DAF (p=0.13, one-sided Wilcoxon signed rank test) **(b)** Mean repeat lengths in the null versus DAF+PS condition. Other plotting conventions as in (a). Panels **(c)** and **(d)** show the same data as panels (a) and (b), respectively, displayed at the absolute difference in mean repeat number. Red dot in (a) and (c) corresponds to the data shown in Figure 6. **(e)** Comparison of the change in mean repeat number induced by the DAF (“delay”) and DAF+PS (“delay + shift”) conditions. No significant difference was detected between the changes in repeat number in the DAF and DAF+PS conditions (p=0.65, one-sided Wilcoxon signed rank test).

## Discussion

Manipulation of auditory feedback induced robust sequence changes in the song of adult Bengalese finches. As hypothesized, both DAF and DAF+PS induced changes in transition probabilities and repeat length distributions in a substantial number of individual cases (filled symbols, Fig. 4a and 7a, b). At branch points, both feedback manipulations induced a reduction in the probability of the primary transition (Fig. 4a, b). In contrast, whereas DAF did not significantly bias changes in mean repeat number upwards or downwards (Fig. 7a, c), DAF+PS caused a reduction in mean repeat number in a significant fraction of cases (Fig. 7b, d). Together, these results demonstrate that continuous, real-time manipulation of auditory feedback can strongly modulate vocal performance and that in some contexts, the addition of a pitch shift can significantly reduce the vocal changes induced by auditory feedback delays.

A number of studies have examined the consequences of subjecting birds to song-triggered playbacks of (previously recorded) samples of the bird’s own song, a manipulation that approximates delayed auditory feedback. In contrast, our technique employs miniaturized headphones to continuously stream manipulated auditory feedback. It is possible that this methodological difference accounts for some apparent discrepancies between our results in the DAF condition and prior findings. Notably, Sakata and Brainard (2006) used playbacks of 1-3 song syllables at specific times during Bengalese finch songs. Similar to our findings, they found that playbacks targeted to branch points reduced the probability of the primary transition (Sakata and Brainard, 2006). In contrast to our findings, however, this earlier paper noted that the effects of feedback manipulation were observed very soon after the manipulation was introduced and did not increase with continued exposure, whereas in many of our experiments (Fig. 3b, 5b) the magnitude of DAF effects on sequence grew steadily over the first few days of exposure. Although the variation in behavioral effects might reflect the different methods of altering auditory feedback, additional studies would be required to isolate the effects of continuous, real-time feedback (our study) versus intermittent, pre-recorded feedback (Sakata and Brainard, 2006) from other methodological differences between the two studies, including the total time of exposure to altered feedback and the magnitude of the feedback delay.

The headphones apparatus greatly attenuates airborne transmission of a bird’s vocalization, replacing it with the manipulated version played through the headphones speakers. However, as discussed elsewhere, subjects might receive unmanipulated acoustic feedback via bone conduction, in which sound is transmitted via body tissues rather than air (Sober and Brainard, 2009). While we cannot rule out some influence of bone conduction, we note that this factor presumably applies in both the DAF and DAF+PS conditions, and therefore seems unlikely to account for the differing effects of these results. We further note that potential bone conduction signals are only one of several sensory modalities that can convey unmanipulated feedback, with proprioceptive/somatosensory systems additionally providing information about the birds’ actual motor output.

Our findings highlight the importance of the characteristics of auditory feedback on vocal behavior. As shown in Figure 4c, DAF elicited significantly larger changes in branch point transition probability than did DAF+PS, as hypothesized. This finding is significant for two reasons. First, it parallels similar findings in persons who stutter. The vocal sequencing errors that typify stuttering can be reduced by the application of an online pitch shift (Kalinowski et al., 1993; Natke et al., 2001; Buchel and Sommer, 2004). Our analogous finding in songbirds (i.e. that delay-induced sequencing changes can be partly reversed by pitch shifts) suggests that songbirds might be used as an animal model of how temporal and acoustic properties of auditory feedback might be manipulated to enhance the fluency of human speech. Second, our findings suggest that pitch shifts reduce songbirds’ reliance on auditory feedback when sequencing vocal behavior. A prior study employing pitch shifts, but not delays, found that while smaller (±0.5 or 1.0 semitone) pitch shifts evoke compensatory changes in vocal pitch, ±3.0 semitone pitch shifts did not evoke robust changes in vocal acoustics (Sober and Brainard, 2012). The present findings suggest that pitch-shifted auditory feedback is similarly disregarded when animals program upcoming vocal sequences.

Our analyses did not reveal any significant difference in the effects on repeat number evoked by DAF and DAF+PS. Although further refinements of our technique, such as testing of other delay magnitudes, might reveal such a difference, it is also possible that these two forms of variable sequencing (branch points and repeated syllables) differ in their reliance on the acoustic structure of auditory feedback (Wittenbach et al., 2015). Future studies could examine this possibility by examining the effects of sensory perturbations on behavior and neural activity during vocal production.

